# Small-molecule ketone esters treat brain network abnormalities in an Alzheimer’s disease mouse model

**DOI:** 10.1101/2022.09.24.509337

**Authors:** John C. Newman, Keran Ma, François Kroll, Erin Higgins, Scott Ulrich, Jorge J. Palop, Eric Verdin

## Abstract

Altered brain network activity and the resulting hypersynchrony are important for the pathogenesis of cognitive decline in Alzheimer’s disease (AD) mouse models. Treatments that reduce epileptiform discharges (EDs) or network hyperactivity improve cognition in AD models and humans. We first show that ketogenic diet, but not fasting, rapidly and persistently reduced EDs in the hAPPJ20 Alzheimer’s mouse model over timescales of hours to months. Then, to identify the specific mechanism of the pleiotropic ketogenic diet, we developed small molecule ketone esters to deliver ketone bodies pharmacologically. Two ketone esters recapitulate ED suppression without other dietary manipulation, over time scales of minutes to one week. This small molecule rescue was associated with reduced low-frequency oscillatory activity similar to the recently reported mechanism of an NMDA receptor modulator molecule in this model. Long-term KD resulted in cognitive improvement and in a sex-stratified analysis also improved survival in the more severely affected hAPPJ20 males. Agents that deliver ketone bodies via small molecules or act on downstream targets may hold therapeutic promise in AD through the mechanism of improved network function and reduced epileptiform activity.

## Introduction

Links between epilepsy and Alzheimer’s disease (AD) are seen in both human patients and mouse models. Human patients with AD may commonly have subclinical epileptiform discharges (EDs) (Vossel et al., 2016), and overt epilepsy is associated with more rapid cognitive decline (Lam et al., 2017; Vossel et al., 2013). Mechanistic studies in mouse models of AD have shown that altered oscillatory activity and EDs stem from dysfunctional inhibitory interneurons (Palop and Mucke, 2016), which are key elements of cortical circuits underlying cognition (Jagirdar and Chin, 2019; Kvitsiani et al., 2013). Pathological tau and Aβ promote network hyperexcitability, including through interactions with other AD risk genes (Kazim et al., 2021; Voskobiynyk et al., 2020). Pharmacological treatments or genetic manipulations that reduce EDs improve cognition in these models, including an NMDA receptor positive allosteric modulator that suppresses epileptiform activity by reducing low-frequency oscillations (Hanson et al., 2020; Martinez-Losa et al., 2018; Merlini et al., 2021; Sanchez et al., 2012; Verret et al., 2012). In humans with mild cognitive impairment (MCI), low doses of the antiepileptic levetiracetam improves network hyperactivity and cognitive performance (Bakker et al., 2012). Thus, targeting subclinical epileptiform activity or network hyperactivity are promising new therapeutic approaches to AD and are now being tested in early clinical trials (Bakker et al., 2015; Bakker et al., 2012; Vossel et al., 2021; Vossel et al., 2017).

Ketogenic diet (KD) has long been used to treat certain forms of epilepsy, though primarily inherited developmental disorders (Keene, 2006). Ketogenic diets are defined by restricting carbohydrate intake sufficiently to induce endogenous synthesis of ketone bodies. Ketone bodies are small molecule metabolites, primarily β-hydroxybutyrate (BHB) and acetoacetate, which are synthesized in the liver from lipids in order to provide an alternative, fat-derived, source of circulating energy when glucose is scarce, such as during fasting or a ketogenic diet. But KD is highly pleiotropic and its mechanism of action varies between epilepsy models, in some models involving glycolysis, insulin, specific fatty acids, or even gut microbiome metabolites rather than ketone bodies (Kim and Rho, 2008; Olson et al., 2018). Aside from clinical use in childhood epilepsies, ketogenic interventions including KD (Lilamand et al., 2021b; Neth et al., 2020; Phillips et al., 2021; Taylor et al., 2018) and medium chain triglyceride supplements (Croteau et al., 2018; Fortier et al., 2021; Lilamand et al., 2021a) are under clinical investigation for AD primarily based on the hypothesis of improved energy metabolism from ketone bodies (Cunnane et al., 2020). However, a wide variety of alternative relevant mechanisms in AD have been proposed, based on the complex physiology of ketogenic diets and the pleotropic molecular actions of ketone bodies (Newman and Verdin, 2017), including inflammatory modulation via NLRP3 (Shippy et al., 2020; Youm et al., 2015), G-protein coupled receptors (Hasan-Olive et al., 2019; Wu et al., 2020), epigenetics (Shimazu et al., 2013), the microbiome (Ang et al., 2020; Ma et al., 2018; Olson *et al.*, 2018), and others.

The key barrier to translation of ketogenic-related therapies for AD is a better understanding of specific relevant mechanisms that can guide more targeted therapies and be linked to translatable biochemical or physiological biomarkers. KD is a challenging intervention to implement in cognitively impaired individuals (Taylor *et al.*, 2018) and the necessarily high fat content can be associated with hyperlipidemia, constipation, electrolyte disturbances and other side effects (Cervenka et al., 2017). Exogenous delivery of ketone bodies is an obvious candidate for a more targeted therapy, if shown to be mechanistically relevant. Yet the ketone bodies BHB and acetoacetate are rapidly-metabolized organic acids, so direct exogenous delivery of pharmacological quantities requires a deleteriously large salt or acid load. Ketone esters are a technology to avoid this difficultly by masking the carboxylic acid moiety in an ester bond with other ketone bodies or ketogenic precursor molecules. The specific design of ketone esters can be targeted to deliver varying ratios of BHB and acetoacetate, to provide only exogenous ketone bodies or stimulate endogenous ketogenesis as well, and to optimize delivery kinetics – once these goals can be defined in a particular therapeutic context.

Here we provide a comprehensive and rigorous multidimensional electroencephalography (EEG) and behavioral study of the concurrent effects of a ketogenic diet or pharmacological delivery of ketone bodies via novel small molecule ketone esters on brain network and epileptiform activity, cognitive decline, and survival, using the well-characterized hAPPJ20 mouse model of AD. We find that a ketogenic diet consistently reduces epileptiform discharges over time frames ranging from minutes to months. To define the specific mechanism, we show that similar reduction of ED is observed using two structurally related ketone esters without ketogenic diet; and that all of these interventions act independent of gamma oscillations associated with inhibitory interneurons but instead the mechanism of ED suppression may be via suppression of low-frequency oscillations. Finally, a ketogenic diet improves context-dependent and visuo-spatial learning in hAPPJ20 mice, and reduces the high seizure-related mortality observed in males. Small molecule therapies derived from ketone bodies may act to ameliorate cognitive dysfunction in AD through reducing subclinical epileptiform activity.

## Results

### Ketogenic diet suppresses AD-related epileptiform discharges (EDs)

hAPPJ20 (APP) mice carry a human APP transgene with the Swedish and Indiana familial AD-causing mutations (J20 line) (Mucke et al., 2000) and display amyloid deposition, cognitive deficits, spontaneous EDs (also known as epileptiform spikes), and early premature mortality (Palop et al., 2007; Verret *et al.*, 2012). In order to test if a ketogenic diet would suppress EDs in hAPPJ20 mice, we designed a pair of diets matched for protein content, fat sources, and micronutrients: a typical high-carbohydrate control diet based on AIN-93M (Control), and a zero-carbohydrate ketogenic diet (KD) (Fig. 1A). In pilot experiments with wild-type (WT) mice, we found that initiation of KD resulted in a sustained increase of blood β-hydroxybutyrate (BHB) levels similar to that of an overnight fast (Fig. 1B), along with reduced blood glucose (Fig. 1C), without reduction in caloric intake (Fig. 1D).

**Figure 1.**
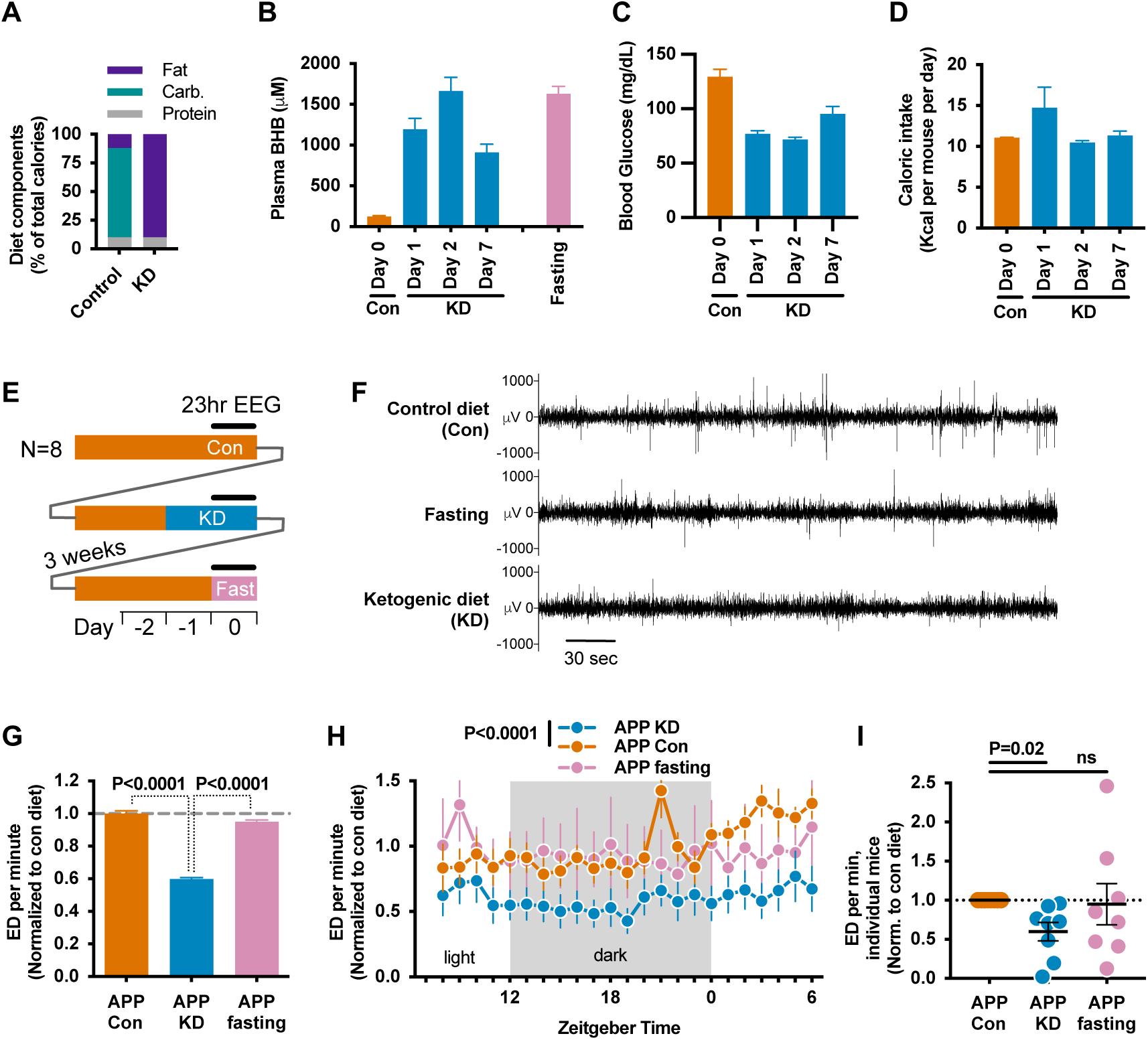
Ketogenic diet reduces epileptiform discharges (ED) in the hAPPJ20 mice. 12-month-old hAPPJ20 mice (N=8: 3M, 5F) were EEG implanted and recorded under control, ketogenic diet, and fasting conditions. **A**, Macronutrient content of control and ketogenic diet (KD). **B**, Initiation of KD increases blood BHB levels similar to a 16-hour fast with a peak at day 2, and **C**, reduces blood glucose. **D**, Initiation of KD does not reduce caloric intake. **E-I**, Longitudinal 23-hour EEG study of KD, fasting, and control diet. **E**, Study schematic. **F**, Representative EEG tracing showing ED. **G**, KD but not fasting reduced ED. **H**, KD reduced ED throughout the 24h circadian cycle. **I**, Response of individual mice to KD or fasting. ED per minute are normalized to control diet EEG recordings for each individual mouse. All data are presented as mean ± SEM. P-values by one-way ANOVA with matching, and Holm-Šídák multiple comparisons test.

We then followed a cohort of hAPPJ20 mice undergoing serial 23-hour EEG recordings under three conditions: on control diet, on the second day after starting KD (to capture peak BHB levels), and during a 23 hour fast (Fig. 1E). Mice recovered on the control diet for 3 weeks between the KD and fasting recordings. We observed that KD, but not fasting, suppressed the rate of EDs (Fig. 1F) by approximately 40% (Fig. 1G). ED suppression was consistently observed throughout the 23-hour recordings on KD (Figs. 1H). ED suppression was also consistent between individual mice on KD, while fasting had more heterogenous effects including exacerbating EDs in some mice (Fig. 1I).

To asses long-term effects of KD on ED suppression, we obtained a series of EEGs as a separate cohort of mice alternated from control diet (baseline) to KD and back to control (washout) over a period of 3 weeks (Fig. 2A). EDs were again rapidly suppressed after starting KD (Figs. 2B and 2C). However, EDs rebounded to at least the prior frequency immediately after returning to the control diet, suggesting a rapidly reversible suppression mechanism. To then test the stability of ED suppression over a longer time frame, we followed two parallel cohorts of hAPPJ20 mice fed control diet or KD for three months, with nine EEG sessions during the latter half of this period (Fig. 2D). This chronic feeding of KD suppressed EDs similarly to the earlier acute feedings throughout this extended period, without any indication of waning or adaptation despite the highly dynamic kinetics observed earlier (Figs. 2E and 2F).

**Figure 2.**
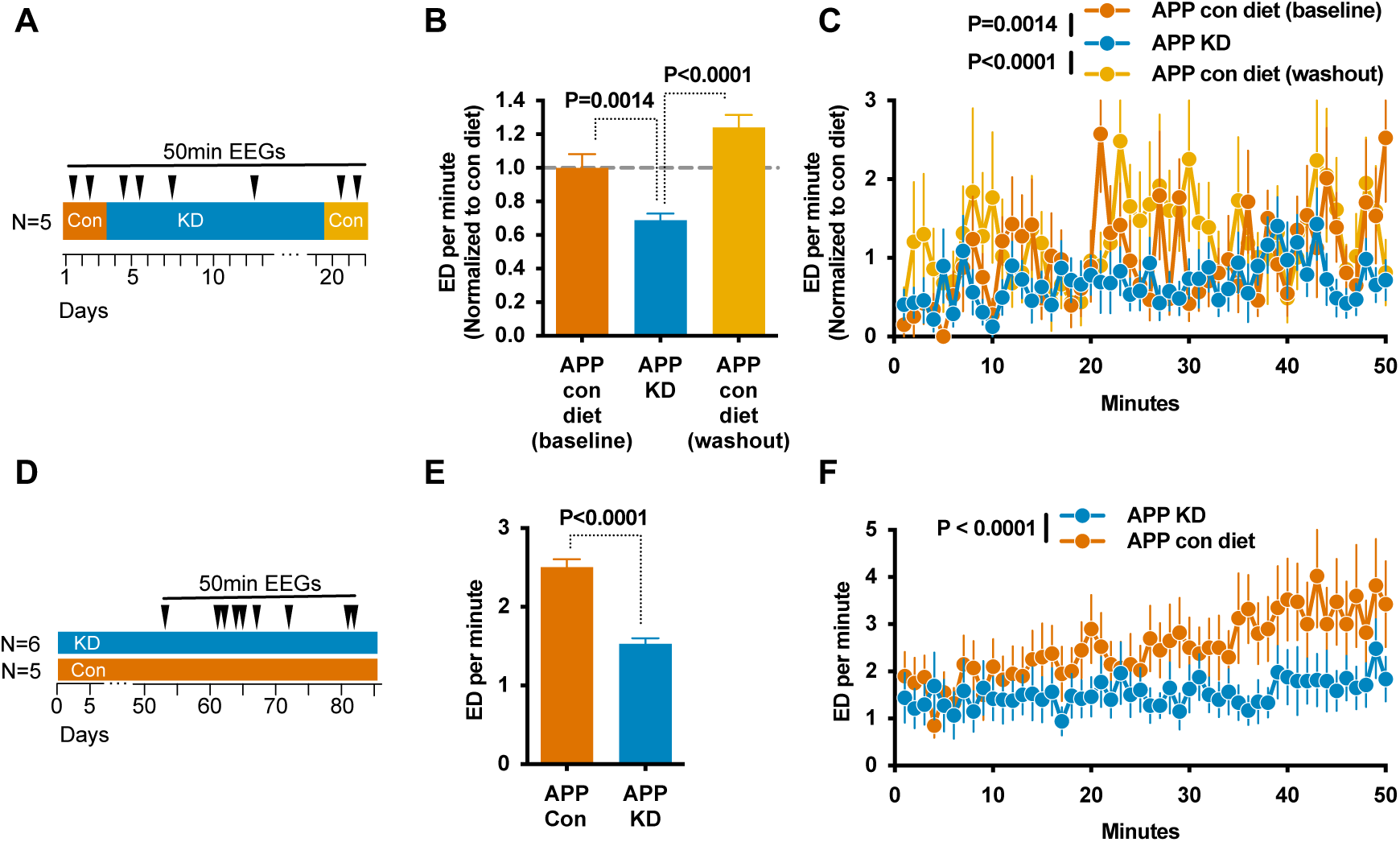
Suppression of epileptiform discharges by ketogenic diet is both dynamic and sustained chronically. **A-C**, Longitudinal cohort of hAPPJ20 mice (10 months old; N=5, 2M, 3F) followed for three weeks onto and off of KD. **A**, Study schematic. KD reduced ED (**B**) throughout the 50 minute recording sessions (**C**). The effect reverted within 2 days of returning to control diet. **D-F**, Parallel groups of of hAPPJ20 mice (8 months old; APP Con *n*=5 (5 M), APP KD *n* =6 (4M, 2F), WT Con *n* =4 (4 M)) were followed on diets for 3 months, with 9 EEG sessions over the last 6 weeks. **D**, Study schematic. Mice on KD again had fewer ED (**E**) throughout the 50 minute recording sessions (**F**). In all panels, data are pooled for all EEGs within a relevant condition. In the longitudinal study (A-C), ED per minute are normalized to the pre-KD control diet EEG recordings for each individual mouse. Normalization is not applicable to the cross-sectional study (D-F). All data are presented as mean ± SEM. P-values by one-way ANOVA with matching (B), two-way ANOVA with (C) or without (F) matching, or T-test (E); with Holm-Šídák multiple comparisons test (B, C).

### Acute and subchronic ED suppression by ketone esters

KD is a complex, pleiotropic intervention with many potential mechanisms for epileptiform spike suppression, including both energetic and signaling functions of BHB, reduced insulin levels, provision of substrates for fatty acid oxidation, and many others (Gavrilovici and Rho, 2021; Newman and Verdin, 2017; Poff et al., 2021; Puchalska and Crawford, 2021). Importantly for potential clinical applications, pleiotropic mechanisms that are not required for ED suppression may underly undesired side effects associated with KD. It is therefore crucial to identify the specific mechanisms of KD that are responsible for ED suppression. A key hallmark of KD is its stimulation of endogenous ketone body production, resulting in circulating blood BHB levels in the millimolar range. To test if ketone bodies are the active component induced by KD in ED suppression, we engineered novel small molecule ketone esters to deliver naturally occurring ketone bodies pharmacologically and therefore bypass the complex diet intervention. In order to best emulate the physiology of ketone body production in KD, we engineered ketone ester molecules that comprise a BHB moiety (for immediate BHB delivery) ester-linked to six-carbon medium chain fatty acids (to stimulate endogenous ketone body production). Medium-chain fatty acids are rapidly metabolized to a physiological and naturally occurring mix of BHB and acetoacetate in the liver regardless of diet context. Compound C6-BHB (Fig. 3A) was selected from other structural derivatives for effectively increasing plasma BHB to fasting/KD-like levels after intraperitoneal injection in WT mice (Fig. 3B) without altering blood glucose (Fig. 3C). In a longitudinal, crossover study of hAPPJ20 mice fed only control diet, we injected both C6-BHB and saline intraperitoneally on separate days and recorded 50-minute EEGs both before and after each injection (Fig. 3D). Injection of C6-BHB increased plasma BHB (Fig. 3E, measured post-recording) and substantially suppressed EDs compared to saline injection (Figs. 3F and 3G). The effect was evident from the first minutes of EEG recording, started 20 minutes after injection. These data are consistent with the model that ketone bodies induced by the ketogenic diet are the key suppressor of EDs, and with the mechanism being dynamic in a time frame of minutes.

**Figure 3.**
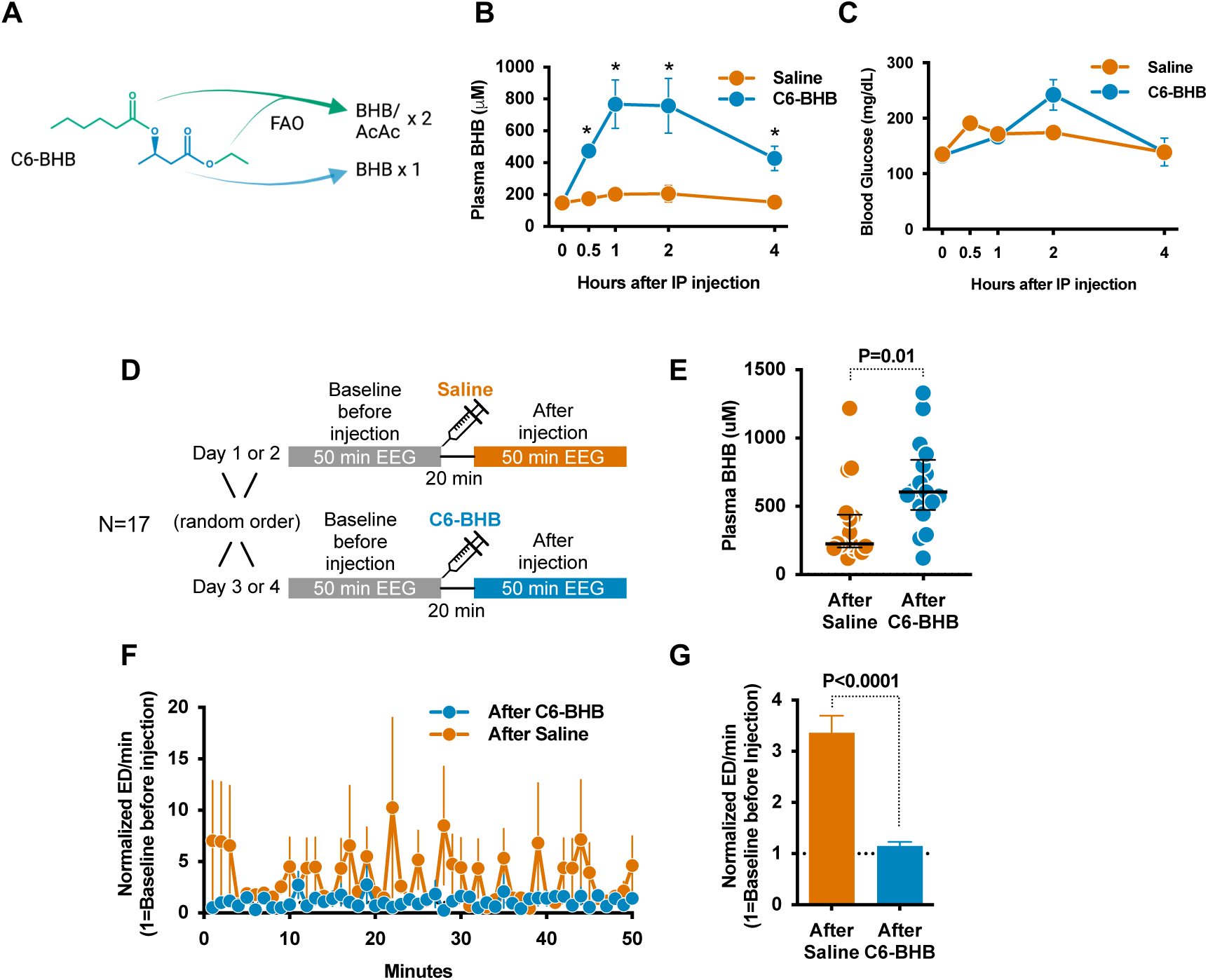
Ketone ester compound injection acutely reduces epileptiform discharges in hAPPJ20 mice. **A,** The C6-BHB ketone ester generates one molecule of BHB directly and 2 molecules of either BHB or acetoacetate via fatty acid oxidation (FAO) of its 6-carbon medium chain fatty acid moiety and ethanol protecting group. **B**, Intraperitoneal injection of C6-BHB in wild-type mice acutely increases blood BHB levels similarly to ketogenic diet, without affecting blood glucose levels (**C**). **D-G**, Longitudinal crossover study of C6-BHB injection vs. saline in hAPPJ20 mice (12-24 months old; N=17, 13M, 4F). **D,** Study schematic. **E**, Increased blood BHB levels 70 minutes after C6-BHB injection. **F**, C6-BHB injection reduces ED throughout the 50 minute recording and in total (**G**) compared to after saline injection. ED per minute are normalized to the pre-injection EEG recordings for each individual mouse injection. All data are presented as mean ± SEM. P-values by paired T-tests (B, G) or Wilcoxon test (E).

In order to asses long-term effects of ketone bodies in ED suppression, independent from diet, we carried out a longitudinal study of administering ketogenic compounds in normal food. We selected two structurally distinct ketogenic compounds, the ketone ester C6×2-BHB (an ester of BHB and medium-chain fatty acids, as described above) and the BHB precursor compound (R/S)-1,3-butandiol which is converted directly to BHB in the liver by dehydrogenases (Fig. 4A). Mice were alternated weekly between control diet and diets containing 10% w/w of a ketogenic compound (Fig. 4B). All diets were carefully matched in macronutrient and micronutrient content except for the substitution of the ketogenic compounds for carbohydrate (Fig. 4C). Both compounds increased plasma BHB (Fig. 4D) without altering blood glucose (Fig. 4E), although 1,3-butanediol resulted in small body weight gain over the 7 days of feeding (Fig. 4F). The ketone ester C6×2-BHB, but not 1,3-butanediol, suppressed EDs similarly to prior KD experiments (Fig. 4G). ED suppression was also consistent between individual mice with C6×2-BHB, while 1,3-butanediol had more heterogenous effects including exacerbating EDs in some mice (Fig. 4H).

**Figure 4.**
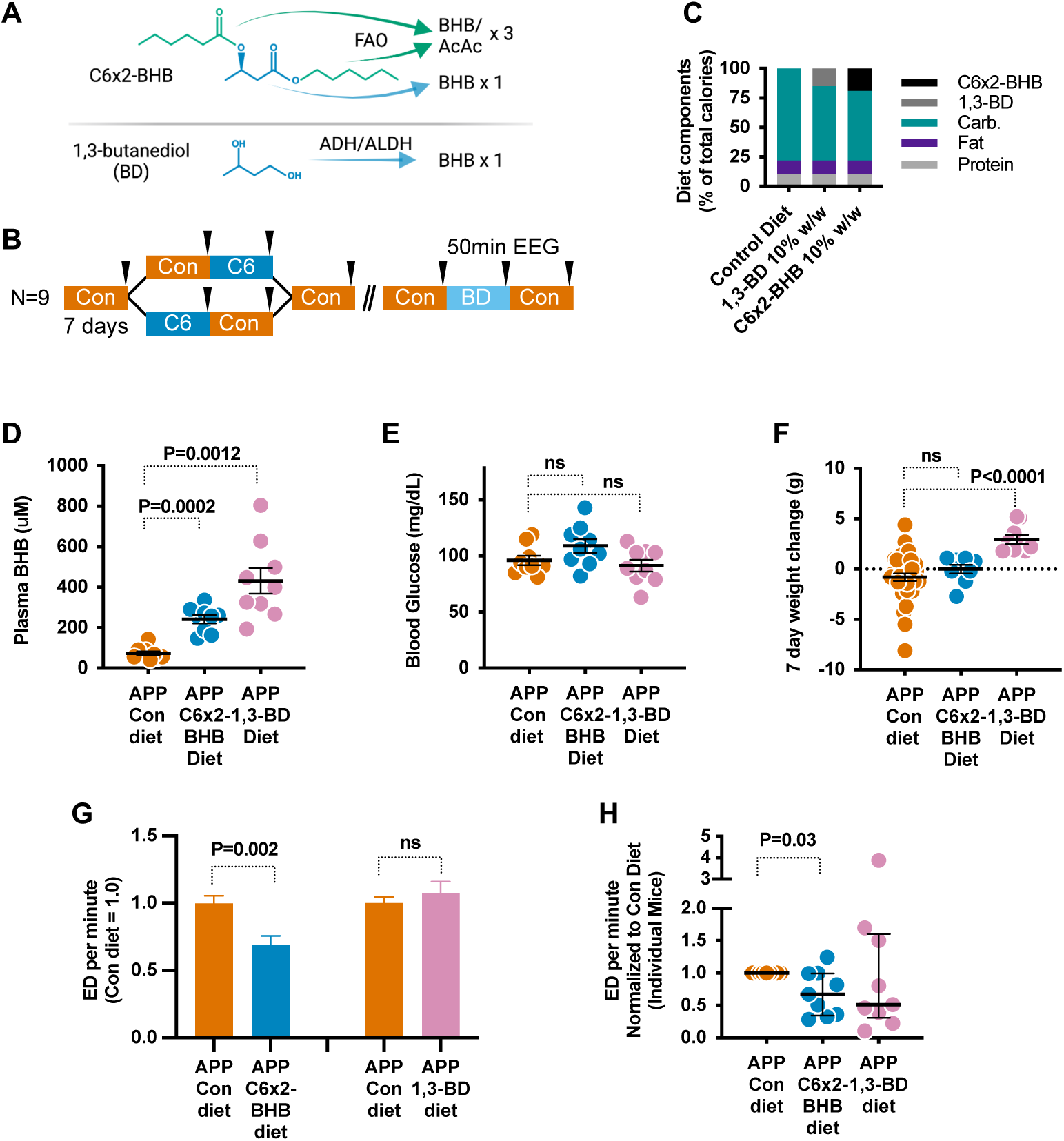
Feeding C6×2-BHB ketone ester, but not 1,3-butanediol (BD), for one week reduces epileptiform discharges in hAPPJ20 mice. **A,** The C6×2-BHB ketone ester, optimized for delivery in food, generates one molecule of BHB directly and 3 molecules of either BHB or acetoacetate via fatty acid oxidation (FAO) of its 6-carbon medium chain fatty acid and medium chain alcohol moieties. **B,** Schematic of the longitudinal study of hAPPJ20 mice (5-14 months old; N=9, 3M, 6F), which included a cross-over feature with an additional control diet period for C6×2-BHB. **C**, Per calorie macronutrient diet composition of the study diets. C6×2-BHB and 1,3-butanediol were provided at 10% w/w. **D**, Both compounds increase blood BHB levels vs. control diet, and have minimal effects on blood glucose (**E**). **F**, Body weight change over the 1 week diet period. **G**, C6×2-BHB, but not 1,3-butanediol, reduces ED compared to control-fed EEGs. **H**, Response of individual mice. ED per minute are normalized to the control diet EEG recordings for each individual mouse during each study period (C6×2-BHB or 1,3-butanediol, as in the schematic). All control diet periods were combined for the metabolic measurements. All data are presented as mean ± SEM. P-values by one-way ANOVA with (D, E) or without (F, G, H) matching and Šídák or Holm-Šídák multiple comparisons test.

### KD and ketone esters do not alter interneuron-related gamma oscillatory activity

EDs in hAPPJ20 mice are caused by the loss of activity of GABAergic interneurons which normally provide a consistent inhibitory tone and oscillatory network activity (gamma activity, measured in the 30–90 Hz gamma frequency range on EEG), thereby preventing pathological hyper-synchronization of neuronal networks (Palop and Mucke, 2010). Interneuron dysfunction in hAPPJ20 mice is associated with loss of expression of the sodium channel subunit Nav1.1 (encoded by Scn1a), predominantly expressed in fasting-spiking interneurons, and cognition is improved by restoring Nav1.1 function (Martinez-Losa *et al.*, 2018; Verret *et al.*, 2012). The antiepileptic drug levetiracetam and the NMDA receptor positive allosteric modulator GNE-0723 have been shown to improve oscillatory network function as well as cognition in AD models (Hanson *et al.*, 2020; Sanchez *et al.*, 2012) In order to test whether KD and ketone ester suppress EDs via enhancing gamma oscillatory activity, we next analyzed the relationships between exploratory movement, interneuron/gamma activity, and EDs beginning with the 23-hour EEG recordings (Fig. 1E). Exploratory movement is associated with increased interneuron-dependent gamma activity and suppression of EDs in hAPPJ20 (Verret *et al.*, 2012). Importantly, neither KD nor fasting altered overall movement relative to control diet (Fig. 5A), indicating that ED suppression from KD was not mediated by KD-induced changes in locomotor activity. Next, we assessed whether gamma oscillations were affected by KD. There was no difference in movement-normalized gamma activity across the 23 hours (Fig. 5B), nor was there a change in the overall fraction of EEG power in the gamma range (Fig. 5C). We used regression analyses to further interrogate the relationships between mouse movement, gamma activity, and EDs. KD reduced EDs at all movement levels (Fig. 5D), and at all levels of movement-adjusted gamma activity (Fig. 5E). Finally, KD did not increase the induction of gamma power with increased movement (Fig. 5F). Similar analysis of the cohort of mice that alternated from control diet to KD and back to control over a period of 3 weeks (Fig. 2A) revealed the same patterns (Supplementary Fig. 1). Overall, these data are consistent with KD acting downstream or independent of gamma oscillations.

**Figure 5.**
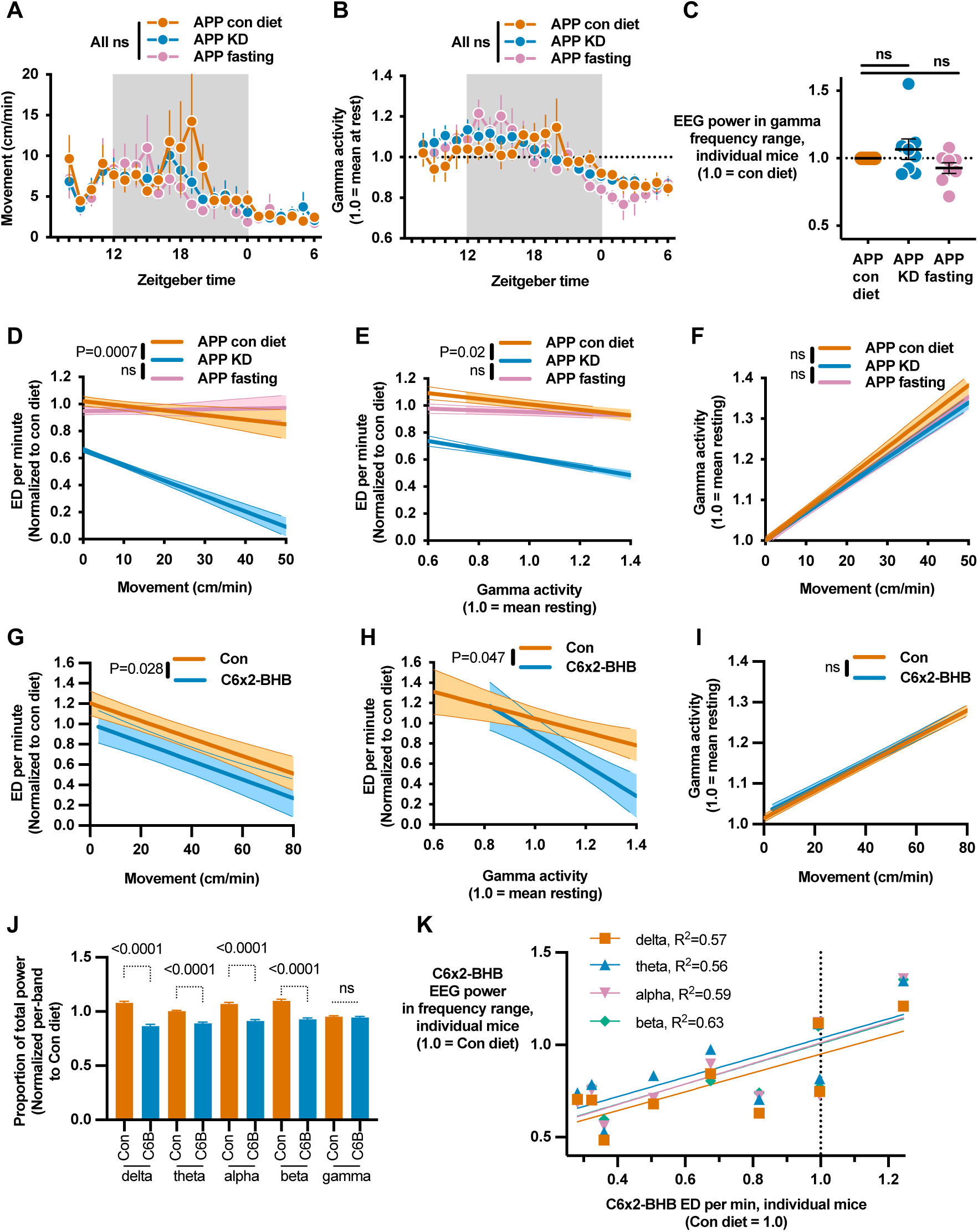
Ketogenic diet and ketone ester do not affect movement or gamma oscillatory activity, but reduce low-frequency oscillations. **A-F,** Data recorded during the 23 hour EEGs in Fig. 1. **A**, Movement was similar in all conditions. **B**, Hourly plot of movement-normalized gamma activity was not different between diet conditions. **C**, The proportion of overall EEG power in the gamma frequency range for individual mice was unchanged by KD or fasting. **D**, Regression plot of per-minute ED versus movement. hAPPJ20 mice on KD exhibited fewer EDs at all movement levels. **E**, Regression plot of ED versus movement-normalized gamma activity showed fewer EDs on KD at all levels of normalized gamma. **F**, Regression plot of movement-normalized gamma versus movement showed no change in the rate of induction of gamma activity by movement for KD or fasting. **G-K**, Data recorded during the C6×2-BHB feeding experiment in Fig. 3. Similar to the KD recordings, mice fed C6×2-BHB show fewer ED at all levels of movement (**G**) and at all levels of normalized gamma (**H**) as well as no change in the induction of gamma activity by movement (**I**). **J**, C6×2-BHB reduces low-frequency power in delta, theta, alpha, and beta bands. **K**, For individual mice, reductions in low-frequency oscillation power is associated with reduced ED. All data are presented as mean ± SEM. P-values for regression lines are computed via ANCOVA for comparison of slopes and then intercepts (D-I), and for other comparisons via one-way ANOVA with matching and Dunnett’s multiple comparison test (C) or no matching and Šídák multiple comparisons test (J).

We found similar results upon analysis of EEG data from the C6×2-BHB ketone ester feeding cohort (Fig. 4). The ketone ester reduced EDs at all movement levels (Fig. 5G) and at all levels of movement-adjusted gamma activity (Fig. 5H), and did not increase the induction of gamma power with increased movement (Fig. 5I). There was no change in the overall fraction of EEG power in the gamma range (Supplementary Fig. 2). C6×2-BHB and 1,3-butanediol both increased movement during the EEGs to a similar degree (Supplementary Fig. 3). A simpler analysis of the smaller injection EEG data set (Fig. 3) similarly showed that ED suppression by C6-BHB was not a result of increasing exploratory movement (mice moved less after C6-BHB than saline injection) and that ED suppression was greater at rest than while moving (Supplementary Fig. 4).

Interestingly, while C6×2-BHB did not alter gamma power, it did reduce power in the lower frequency bands delta, theta, beta, and alpha, spanning 0.5-20 Hz (Fig. 5J). For individual mice, greater power reduction in these bands by C6×2-BHB was associated with greater suppression of EDs (Fig. 5K). Recently, we reported a positive allosteric NMDA receptor modulator that improved cognitive function and reduced epileptiform discharges in hAPPJ20 mice via the mechanism of reducing aberrant low-frequency oscillatory power (Hanson *et al.*, 2020). C6×2-BHB may act via a similar mechanism. Altogether, the ketone esters C6×2-BHB and C6-BHB alone are sufficient to elicit the benefits of KD and appear to act via a similar or identical mechanism to ketogenic diet, suppressing EDs independently of gamma oscillations possibly through rescue of aberrant low-frequency oscillations, and these effects are not explained by alterations in locomotor activity.

### KD improves long-term memory and survival in hAPPJ20 mice

Finally, we investigated whether ED suppression by KD is associated with long-term improvements in memory or survival in hAPPJ20 mice. These mice display characteristic deficits in context-dependent learning, which can be observed as a failure to habituate to a novel environment. During a three-month study (Fig. 2D), mice were exposed to the open field four times in the first month of treatment as training, followed a month later by two test open field sessions to assess if familiarity with the open field from the prior training would reduce exploratory activity (Fig. 6A). During the test open field sessions, hAPPJ20 mice on the control diet exhibited hyperactivity and high levels of exploratory movements, as measured by total movement, center movement and rearings (Fig. 6B-D), demonstrating a lack of habituation that reflects impaired memory. Wild-type (WT) mice showed strong habituation, with low exploratory movement in the test open fields. Strikingly, hAPPJ20 mice on KD displayed habituation as strong as WT mice, with reduced total movement (Fig. 6B), and reduced exploratory movements such as movements through the center of the open field (Fig. 6C) and rearings (Fig. 6D).

**Figure 6.**
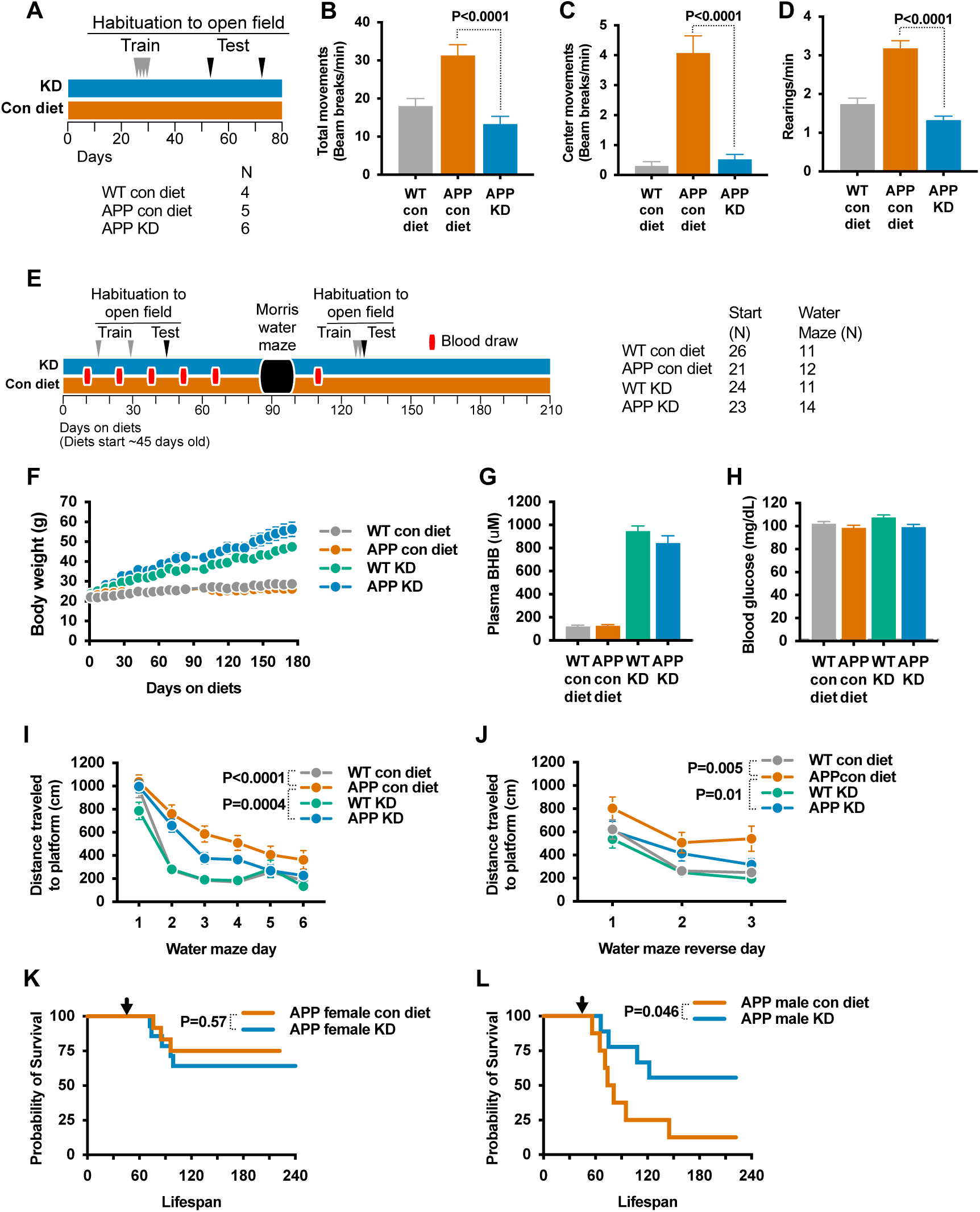
Ketogenic diet improves memory and reduces mortality in hAPPJ20 mice. **A-D,** habituation to open field context-dependent learning test on the same cohort from Fig. 2D. **A**, Study schematic. During the test open fields, APP mice on KD showed normal levels of movement (**B**), center movement (**C**), and rearings (**D**), indicating successful learning. Data from the two test open field sessions are pooled. **E-L**, 7-month study of KD in hAPPJ20 mice starting at 6 weeks old. At start of study, APP Con *n*=21 (9M, 12F), APP KD *n*=23 (9M, 14F), WT Con *n*=26 (12M, 14F), WT KD *n*=23 (16M, 7F). For water maze, APP Con *n*=12 (3M, 9F), APP KD *n*=14 (5M, 10F), WT Con *n*=11 (6M, 5F), WT KD *n*=11 (6M, 5F). **E**, Study schematic. Mice gained weight on KD (**F**), had persistently elevated blood BHB (**G**), and normal blood glucose (**H**). KD improved visuo-spatial learning performance for APP mice during both forward **(I)** and reverse **(J)** training in the Morris water maze. KD did not affect survival of female hAPPJ20 mice **(K)** but markedly improved male survival (**L**). Arrows show the start of study diets (45 days old). All data are presented as mean ± SEM. P-values via one-way (B-D) or two-way (I, J) ANOVA with Tukey’s multiple comparisons test, or log rank survival test (K, L).

To further investigate the effect of KD on memory and premature mortality, we undertook a larger, 7-month study, beginning with 2-month-old hAPPJ20 and WT mice (Fig. 6E). KD was associated with significant weight gain in both genotypes (Fig. 6F) and higher mean daily calorie intake (Supplementary Fig. 5). Six blood draws obtained every ∼2 weeks from the start of the study showed that KD increases plasma BHB levels, averaging ∼1 mM over the 6-month period which is ∼10-fold higher than controls (Fig. 6G). Blood glucose levels were similar in all groups (Fig. 6H). Two separate studies of habituation to the open field completed in month 2 (Supplementary Fig. 6A-C) and month 5 (Supplementary Fig. 6D-F) again showed reduced exploratory center movement and rearings for hAPPJ20 mice on KD in comparison to the control diet, indicating enhanced context-dependent learning. Importantly, analysis of the first open field from both the earlier cohort (Supplementary Fig. 7A-B) and this cohort (Supplementary Fig. 7C-D) gave no suggestion of an anxiolytic effect of KD (Kashiwaya et al., 2013).

During Morris water maze testing, hAPPJ20 mice on KD showed significantly better performance in the hidden-platform training (learning) trials of the water maze than hAPPJ20 mice on the control diet (Fig. 6I). This improvement remained consistent when the location of the platform was moved during reversal training (Fig. 6J). However, there was no difference in performance during the probe (memory) trial of the water maze, either after initial hidden platform training or after reversal training (Supplementary Fig. 8). There was no clear association between sex and performance in either habituation to the open field or water maze (Supplementary Fig. 9). hAPPJ20 mice exhibit substantial early mortality, particularly in males(Davis et al., 2020; Verret *et al.*, 2012) that may be due to fatal seizures. KD did not affect the relatively lower mortality among females (Fig. 6K). Remarkably, however, the high early mortality among males was strongly reduced (Fig. 6L).

Reports of the effect of KD or ketone bodies on Aβ deposition have been conflicting, with Aβ either reduced (Kashiwaya *et al.*, 2013; Van der Auwera et al., 2005; Xu et al., 2022) or unchanged (Aso et al., 2013; Beckett et al., 2013; Brownlow et al., 2013). While changes in plaque accumulation could not explain the very rapid kinetics we observed for the effect of KD and ketone esters on epileptiform activity, we nevertheless carried out an exploratory immunohistochemical analysis of mice after 7 months on KD (Fig. 6E) that revealed no obvious difference in the number or size of Aβ plaques (Supplementary Fig. 10A-B). Ectopic expression of neuropeptide Y (NPY) increases with aberrant network hyperexcitability in hAPPJ20 (Palop et al., 2011a). We indeed observed increased NPY expression in APP mice on the control diet, while expression in APP mice fed KD trended towards WT levels (Supplementary Fig. 10C) consistent with long-term improvement in neuronal hyperexcitability.

## Discussion

In summary, we found that KD suppresses epileptiform discharges, improves context-dependent learning and visuo-spatial memory performance, and increases survival in the hAPPJ20 mouse model of AD. ED suppression can be attained with similar effect size and electrophysiological profile by small molecule ketone ester compounds that provide ketone bodies pharmacologically. These data provide a mechanistic understanding of earlier observations of visuo-spatial memory improvement with KD (Xu *et al.*, 2022; Yin et al., 2016) or a ketone ester diet (Kashiwaya *et al.*, 2013) in AD models. We are the first to show that KD reduces AD-induced epileptiform activity, providing a functional mechanism of action linked to rapid cognitive decline in AD patients (Lam et al., 2017; Vossel et al., 2013; Vossel et al., 2016) and with translational applicability in patients (Bakker et al., 2015; Bakker et al., 2012; Vossel et al., 2021; Vossel et al., 2017). The magnitude of ED suppression is similar to that obtained by low doses of the antiepileptic drug levetiracetam (Sanchez *et al.*, 2012) or by transgenic expression of SCN1A in interneurons (Verret *et al.*, 2012). We further show for the first time that ketone bodies are the functional component of KD for this mechanism, and a small molecule ketone ester alone can fully replicate ED suppression. We carefully define the kinetics of ED suppression by KD and ketone esters, and we identify a relevant potential intermediate mechanism in reduced low-frequency oscillatory activity. A key strength of this study is its scope and replicability, with a large number of mice of both sexes and varying ages analyzed in 6 separate cohorts testing different timing, intervention, and behaviors, all with consistent results supporting the rigor of the conclusions.

Our data argue strongly that ketone bodies are the relevant mechanism of ED suppression in this model, given the close similarity in effect size and EEG characteristics between the KD and ketone ester cohorts. The downstream mechanism of ketone bodies in ED suppression is likely multifactorial, but the options are constrained by our observed kinetics. We found the rescue to be robust, immediate, and stable over time. Direct actions on neurotransmitter signal transduction could fit these kinetics. The detailed EEG analysis argues against modulation of inhibitory interneuron-associated gamma oscillations, but interestingly the ketone ester suppressed low frequency oscillatory activity similar to our recent report of ED suppression by the NMDA receptor positive allosteric modulator GNE-0723 (Hanson *et al.*, 2020). Low-frequency oscillations are associated with the brain resting state and network hypersynchrony in AD models, while suppression in this band favors an active brain state and desynchronized neuronal activity. GNE-0723 acts specifically on GluN2A-subunit-containing NMDRs to enhance receptor activation by endogenous glutamate release. Ketone bodies have been implicated in glutaminergic signaling, BHB as a potential antagonist of GluN2A-subunit containing NMDRs (Pflanz et al., 2019) and acetoacetate as an suppressor of synaptic glutamate reuptake via VGLUT (Juge et al., 2010). While these do not explain our findings, together they may point to an important new activity of ketone bodies in modulating low frequency oscillatory activity via a glutamate-centric mechanism.

Changes in Aβ accumulation, mitochondrial biogenesis, inhibitory interneuron survival, or chronic inflammation, all of which have slow onset and would have increasing effects over time, are unlikely to explain these kinetics. Of course, these other mechanisms, even if not required for immediate ED suppression, may affect network function and/or memory function over longer time frames, as might also the reported increased survival of interneurons with ketone ester feeding (Cheng et al., 2020), or improved neurovascular function with ketogenesis (Lin et al., 2015; Ma *et al.*, 2018).

Our findings are consistent with a recent report of Sirt3 haploinsufficiency exacerbating EDs and mortality in APP/PS1 mice (Cheng *et al.*, 2020), as Sirt3 is an important regulator of ketogenesis (Rardin et al., 2013; Shimazu et al., 2010). A key goal of future studies will be to parse the roles of the two major ketone bodies BHB and acetoacetate. While BHB is the predominant ketone body generated both in KD and from the ketone esters, these all produce acetoacetate as well at a physiological ratio. It is surprising that 1,3-butanediol, which alone among the interventions exclusively produces BHB, generated inconsistent EP spike suppression. However, (R/S)-1,3-butanediol produces both the endogenous R-enantiomer of BHB as well as the non-physiologic S-enantiomer, and while they share many common molecular actions (Newman and Verdin, 2017), it is possible S-BHB acts counter to R-BHB in this context. The inconsistent efficacy of fasting, being a potent inducer of ketone bodies, is also surprising and may be due to a counteracting mechanism or exacerbating factor in the complex metabolic milieu of fasting. An additional difference is that KD and both ketone esters provide substrates for endogenous ketogenesis, while 1,3-butanediol does not. The bulk of ketogenesis occurs in the liver with resulting systemic circulation of BHB and acetoacetate, and from the brain’s perspective there may be no difference between exogenous and liver-derived ketone bodies. But there is growing evidence for local and paracrine roles for ketogenesis. For example, ketone bodies act locally in the liver to regulate fibrosis (Puchalska et al., 2019), and extrahepatic ketogenesis in the gut has important functions in regulating stem cells (Cheng et al., 2019) and perhaps immunity (Ang *et al.*, 2020). There may similarly be a relevant role in network function for local ketogenesis in the brain (Guzmán and Blázquez, 2001; Le Foll and Levin, 2016), which would be stimulated by KD and the C6-BHB/C6×2-BHB ketone esters, but not by 1,3-butanediol.

How KD and ketone bodies affect synaptic function is also an important area for future study in light of our EEG findings. Relevant molecular mechanisms might include modulation of potassium channel activity (Ma et al., 2007), deacetylase inhibition (Min et al., 2015; Shimazu *et al.*, 2013), protein β-hydroxybutyrylation (Xie et al., 2016), or increased GABA synthesis (Roy et al., 2015; Yudkoff et al., 2007). Interestingly, one recent study found that a GABA receptor agonist acutely reduced EDs in Sirt3^+/-^ APP/PS1 mice (Cheng *et al.*, 2020), while another reported that the mechanism of KD in a different mouse epilepsy model was via gut-derived gamma-glutamylated amino acids altering brain GABA levels (Olson *et al.*, 2018). However, there is no evidence that gabapentin or pregabalin are useful in Alzheimer’s disease, and both are noted in the American Geriatrics Society’s list of potentially inappropriate medications for the elderly due to interactions with other drugs and renal function (American Geriatrics Society 2019 Updated AGS Beers Criteria® for Potentially Inappropriate Medication Use in Older Adults, 2019).

While caution is appropriate in considering complex dietary or novel pharmacological interventions for elderly patients with dementia, compounds that enhance ketogenesis or provide ketone bodies pharmacologically such as ketone esters (Newport et al., 2015), or that recapitulate the downstream molecular mechanisms of ketone bodies as these are elucidated in deepening levels of detail, hold promise as therapies for AD. In addition, EEG monitoring of epileptiform activity could provide an important biomarker for efficacy of such agents in clinical trials.

## Acknowledgements

We thank Brett Mensh for advice and comments on the manuscript, Giovanni Maki for assistance with figure graphics, and Gary Howard for editorial assistance. Behavioral data were obtained with the help of the Gladstone Institutes’ Neurobehavioral Core (supported by US NIH grant P30NS065780). This work was supported by US NIH grants K08AG048354 and R01AG67333 (J.C.N.), and RF1AG062234, R01AG062629, P01AG073082 (J.J.P.); Gladstone and Buck intramural funds (E.V.); funds from the Larry L. Hillblom Foundation; and Alzheimer’s Association grant IIRG-13-284779 (J.J.P.). Some figure illustrations were created with BioRender.com. All data needed to evaluate the conclusions in the paper are present in the paper and/or the Supplementary Materials. Additional data can be provided by the authors pending scientific review and a completed material transfer agreement. Data requests should be submitted to the corresponding author.

## Authorship

J.C.N., J.J.P. and E.V. conceived the studies and designed the experiments. J.C.N., K.M., F.K., and E.H. carried out the experiments. S.U. and E.H. synthesized the novel small molecules. J.C.N. performed data analysis and wrote the manuscript. J.J.P. and E.V. provided supervision, training, and mentorship at all stages.

## Methods

### Animal Care

All mice were maintained according to the National Institutes of Health guidelines, and all experimental protocols were approved by the University of California San Francisco (UCSF) Institutional Animal Care and Use Committee (IACUC). UCSF is accredited by the Association for Assessment and Accreditation of Laboratory Animal Care (AAALAC). Mice were maintained in a barrier facility on a 7:00 am to 7:00 pm light cycle and had free access to water. Except if stated otherwise, they were housed in littermate groups of up to 5 mice per cage, and fed ad libitum a standard chow diet (5053 PicoLab diet, Ralston Purina Company, St. Louis, MO).

### Mouse strains

hAPPJ20 mice carry a transgene containing human APP with the Swedish and Indiana FAD mutations, on a C57BL/6J background (Mucke *et al.*, 2000). Wild-type controls are littermates that do not carry the APP transgene. Some ancillary experiments involving effects of ketogenic diet or novel compounds were performed using wild-type C57BL/6 male mice from the National Institute on Aging Aged Rodent Colony, usually obtained at 11 months of age, with experiments carried out between 11 and 16 months of age.

These mice were tested and found to carry the Nnt partial gene deletion common to certain C57BL/6 substrains.

### Experimental diets

Customized ketogenic and control diets were obtained from Envigo. The control diet is based on AIN-93M, and contains per-calorie 10% protein, 13% fat, and 77% carbohydrates (TD.150345). The ketogenic diet contains per-calorie 10% protein and 90% fat (TD.150348). The primary fat sources are Crisco and corn oil in both diets. Fatty acids in the ketogenic diet are, by weight, approximately 24% saturated, 39% monounsaturated, and 37% polyunsaturated. The ketogenic diet is matched to the control diet on a per-calorie basis for micronutrient content, fiber, and preservatives. The compound diets contained 10% w/w of either the ketone ester C6×2-BHB (∼8 kcal/g) or 1,3-butanediol (∼6 kcal/g). They were otherwise also matched to the control diet on a per-calorie basis for protein content (10% of kcal), fat content (12% of kcal), micronutrients, fiber, and preservatives. Note that the standard vivarium chow (PicoLab 5053) contains per-calorie 24% protein, 13% fat, and 62% carbohydrates. The ketogenic diet is of dough-like texture that permits it to be placed in the food well of the cage-top wire lid, in the same manner as pellets. All other diets are firm pellets. All diet were fed *ad libitum* to avoid confounding effects from partial fasting, with caloric intake and body weights tracked. All custom diets were changed weekly.

### Mouse cohort descriptions

In all figures, “APP” is hAPPJ20; “WT” are wild-type littermates that do not carry the hAPP transgene; “Con” or “con diet” is the control diet; “KD” is ketogenic diet.

Figure 1E-I: *N*=8 (3M, 5F), all hAPPJ20.

Figure 2A-C: *N=5* (2M, 3F), all hAPPJ20

Figures 2D-F and 6A-D: APP Con *n*=5 (5 M), APP KD *n* =6 (4M, 2F), WT Con *n* =4 (4 M).

Figure 3D-G: *N*=17 (13M, 4F), all hAPPJ20.

Figure 4B-H: *N*=9 (3M, 6F), all hAPPJ20.

Figure 6E-L: At start of study, APP Con *n*=21 (9M, 12F), APP KD *n*=23 (9M, 14F), WT Con *n*=26 (12M, 14F), WT KD *n*=23 (16M, 7F). For water maze, APP Con *n*=12 (3M, 9F), APP KD *n*=14 (5M, 10F), WT Con *n*=11 (6M, 5F), WT KD *n*=11 (6M, 5F). For immunohistochemistry (Figure 6 – supplement 6), APP Con *n*=2 (1M, 1F), APP KD *n*=5 (3M, 2F), WT Con *n*=3 (1M, 2F), WT KD *n*=4 (2M, 2F).

### Blood draws

Blood for plasma BHB testing was obtained via minimal distal tail snip, with mice placed into a whole-body restrainer (Braintree Scientific) for comfort. ∼40 µL whole blood was drawn for a BHB assay, and collected into microvettes coated with Li-Heparin (Sarstedt). Plasma was separated by centrifugation at 1500 x G for 15 min at 4 °C, and kept frozen at – 20 °C until use. Unless noted, blood draws were done in the morning shortly after lights-on (8-11am).

### Blood assays

BHB concentrations were determined from plasma using a BHB enzymatic detection kit (Stanbio Laboratory, Boerne, TX). Reactions were run with 3 uL of plasma each, in triplicate. Background absorbance recorded before substrate addition was subtracted form the final absorbance to correct for any hemolysis in the plasma samples.

Absorbance was compared to a standard curve. We validated that a standard curve made using the kit-supplied standard (0-1 mM BHB) provided linear values up to 4 mM by testing 1-10 mM BHB solutions prepared fresh from powder (Sigma 298360). Blood glucose was evaluated on a drop of whole blood using a commercial glucometer (FreeStyle Freedom Lite).

### Electroencephalograms

hAPPJ20 mice were implanted (under anesthesia) with Teflon-coated silver wire electrodes (0.005-inch diameter) attached to a microminiature connector bilaterally into the subdural space over frontal, central, parietal, and occipital cortices. For EEG recordings, mice were placed into clear plastic containers that were either 50 cm diameter cylinders (injection experiments) or 20×20 cm boxes (all other non-open-field recordings). Mice are free to move around the container during recordings. For 23-hour recordings, the container was lined with absorbent bedding and mice were provided with food in a small glass jar and water via gel-pack. For other recordings, the floor of the container was bare. Some EEGs were recorded during open-field experiments; this apparatus is described below. All apparatuses were disinfected with Vimoba prior to use and cleaned with 70% ethanol between mice.

EEG recording was performed with Harmonie software from Stellate Systems. Gotman spike detectors from Harmonie were used to automatically detect EDs, which is defined by a peak lasting less than 80 ms and reaching an amplitude greater than five-fold the average baseline amplitude measured during the 5 sec before the spike (Gheyara et al., 2014; Palop *et al.*, 2007). Randomly selected subsets of recordings were reviewed manually to confirm validity of the automated spike detection. The algorithm overcounted a mean of 0.015 EDs/min (median 0.013) in recordings where zero EDs were manually identified, and undercounted by a mean of 9% (median 0%) in recordings where any EDs were manually identified. The number of EDs detected were tallied over each sequential 1-minute window of recording. In all longitudinal experiments, EDs are reported as normalized to the control condition for each individual mouse. This compensates for the variability of baseline EDs between individual mice, and ensures that each mouse is equally represented in the statistical analysis. In other words, it ensures that effect sizes and statistical testing are not unduly influenced by any individual mouse with disproportionately large numbers of EDs at baseline, or by an unusual variance in automated spike detection at very high spike rates in an individual mouse. For parallel-group experiments for which mice are subject to only one condition each (Fig. 2d), ED totals are reported as raw counts.

Mice were also videotracked during EEG recordings, and their movement quantified using Noldus Ethovision software. Videotracking during dark hours (23-hour recordings) was accomplished with infrared illuminators. Ethovision generated raw position coordinates 25-33 times per second. Despite optimization of the tracking parameters, overnight recordings in particular contained artifactual movement from centerpoint jitter and from transient aberrant detection of shadows or reflections. After extensive analysis of the movement data, we arrived at the following custom algorithm to remove artifacts: 1. Calculate a running median of raw positions across 1 second of data intervals; 2. Using the median positions, calculate the distance moved for each data interval (1/25-33 sec); 3. Ignore the distance if it fails any of three criteria: A) velocity >125cm/sec, B) direction of movement not within 90 degrees of the prior interval, C) increase in velocity >8-fold from prior interval. Although overnight (dark) recordings had the most artifacts requiring cleaning, this algorithm was applied to all movement data from all EEG experiments.

Raw power in various frequency ranges was quantified from the EEG recordings using ADInstruments LabChart software. Frequency ranges were defined as follows: 0.5-4 Hz (Delta), 4-10 Hz (Theta), 10-13 Hz (Alpha), 13-20 Hz (Beta), 30-58 Hz and 62-90 Hz (Gamma). 58-62 Hz was excluded to avoid artifacts from the 60 Hz electrical current. Power was calculated in 0.5 sec intervals, and the median of such intervals taken over each discrete minute. This per-minute power was then related to per-minute movement and EDs. The absolute magnitude of raw power is affected by various incidental factors such as length of wire, strength of wire connections, etc., and so cannot be compared directly between mice. Instead we calculated two normalized measures of Gamma frequency power: Adjusted Gamma and Gamma Fraction. Gamma Fraction is defined as the total power in the Gamma range as a fraction of the total power in all specified frequency ranges (Delta, Theta, Alpha, Beta, and Gamma). Adjusted Gamma is normalized so that 1.0 is the mean level of gamma power when the mouse is not moving. For each individual recording session (one mouse, one recording) we performed a linear regression of the set of all per-minute movement and gamma data. 1.0 is defined as the Y-intercept of the regression line, i.e. the gamma power predicted by the regression at zero mouse movement. All gamma power data points are then normalized as a simple ratio to this Y-intercept (Adjusted Gamma = [Raw Gamma] / [Regression Gamma Value at Y-Intercept]).

The several data types from the EEG experiments (EDs, movement, and frequency power) were time-correlated for regression analysis using custom software.

### Open Field-EEGs and Habituation to the Open Field

The open field apparatus (automated Flex-Field/Open-Field Photobeam Activity System; San Diego Instruments) consisted of four identical clear plastic chambers (40 x 40 x 30 cm) with two 16 x 16 photobeam arrays to detect horizontal and vertical (rearing) movements. Total movements were reported, and then stratified as peripheral (4 beams on either flank of each array; 8 beams total) or central (middle 8 beams of each array). Habituation to the Open Field involves exposing mice serially to the open field environment, over which time normal/wild-type mice demonstrate reduced movement and exploratory behaviors. For EEG Open Fields, mice were prepared for EEG as above, and EEG data was collected with Harmonie software from Stellate Systems as above. Data analysis for EDs and raw power were as above. Mice were not videotracked in the Open Field, but movement data was gathered from beam breaks.

### Morris Water Maze

The water maze pool (122 cm diameter) contained water made opaque with powdered white paint, with a 14 x 14 cm platform submerged 2 cm below the surface. For spatial training sessions, mice were trained to locate the hidden platform over at least 5 consecutive days (two sessions of two trials per day, 4 hours apart). The platform location was constant in hidden platform sessions and entry points were changed semirandomly between trials. Mice that failed to find the platform within the time limit of a training trial were placed briefly on the platform before being removed from the pool. The platform was removed for the 60 second probe trials, performed 4 hrs (followed by re-training), 24 hrs, and 48 hrs after the final training session. 24 hr probe trial data is presented. After the conclusion of the initial training and probe trials, the platform location was moved for reverse training, which involved two days of additional hidden platform training on the new location (two trials/day) followed each day 4 hrs after training by a probe trial with the platform removed. The final probe trial (after four total training trials) is presented. To control for vision and motor performance, cued training sessions with a black and white striped mast mounted above the platform were performed later. Mice were videotracked and their movement quantified during the water maze sessions with Ethovision software (Noldus).

### Ketone Ester Compounds

The ketone ester compounds C6-BHB and C6×2-BHB were synthesized by Scott Ulrich of Ithaca College (Ithaca, New York, USA). All compounds were synthesized using the R-enantiomer of BHB as starting material, and they retain this chirality in the ester form. The common names of the compounds are: hexanoyl ethyl β-hydroxybutyrate (C6-BHB) and hexanoyl hexyl β-hydroxybutyrate (C6×2-BHB). These acyl substituted ethyl β-hydroxybutyrate esters used ethyl β-hydroxybutyrate as a starting material, dissolved in pyridine and reacted with hexyl or octyl acyl chloride. The product is diluted with ethyl acetate, and washed with hydrochloric acid, sodium bicarbonate, and brine. The ethyl acetate layer is removed via drying with magnesium sulfate; the pyridine solvent is removed by rotary evaporation and vacuum pumping. ^1^H NMR and gas chromatograph mass spectrometry confirmed >95% purity of the final product. Hexyl β-hydroxybutyrate was prepared by suspending sodium β -hydroxybutyrate in dry dimethylformamide and reacting with 1-bromohexane. Washing and purification proceeded as above, except that the acid wash steps were omitted, and the purity confirmed as above before proceeding to the reaction with hexyl acyl chloride.

The ketone ester compounds are rapidly hydrolyzed upon ingestion by estereases that are ubiquitous in the gut and in blood, with little or no intact compound detectable circulating in blood (Stubbs et al., 2021). For C6-BHB, hydrolysis releases one molecule each of BHB, hexanoic acid, and ethanol. For C6×2-BHB, hydrolysis releases one molecules of BHB and two molecules of hexanoic acid. Hexanoic acid is a medium-chain fatty acid that is rapidly metabolized to ketone bodies (BHB and acetoacetate in a physiological ratio) in the liver regardless of dietary context.

### Injection-EEGs

hAPPJ20 were prepared for EEG as above. Mice underwent a 50 minute baseline EEG recording, followed immediately by intraperitoneal injection. They were allowed to rest 20 minutes in the home cage, then underwent a second, post-injection 50 minute EEG recording. All mice were tested with both compound and saline injections, on different days. The order of injecting compound and saline was randomized. Mice were tested in the same order on each day, so the time of day would be similar for both compound and saline injections for each mouse. Intraperitoneal injections were of 50 µL of pure compound or saline (150 mM sterile sodium chloride in water); for C6-BHB, this volume represented 0.2 millimoles of compound. Blood was drawn immediately after the post-injection EEG, or approximately 70-80 minutes after injection. Data analysis was limited to mice that completed both injections (compound and saline) and all four EEG recordings. hAPPJ20 mice were maintained on custom control diet for two weeks before and during the experiment. Note that most mice showed an increase in EDs after saline injection, due to the stress of the procedure.

### Immunohistochemistry

Exploratory analyses of Aβ plaques and NPY expression were carried out on a subset of mice randomly seleted from the cohort described in Fig. 6E-L. Other mice from this cohort were either euthanized or maintained for other exploratory studies. For Aβ plaques and NPY immunohistochemistry, mice were transcardially perfused with phosphate buffered saline (PBS) and hemibrains were drop-fixed in 4% paraformaldehyde in PBS at 4°C for 24h and cryoprotected in 30% sucrose in PBS (Verret *et al.*, 2012). Brains were coronally sectioned with a sliding microtome into 30-um floating sections. Sections were stored in a solution consisting of 30% glycerol, 30% ethylene glycol, and 40% PBS at –20°C until staining. Endogenous peroxidases were blocked with 3% hydrogen peroxide and 10% methanol in PBS and nonspecific binding with 10% normal donkey serum, 2% gelatin, and 0.5% Triton in PBS. Primary antibodies used were biotinylated 82E1 (1:1000; Immuno-Biological Laboratories) for Aβ plaques detection and rabbit anti-NPY (1:2000; Immunostar). NPY Primary antibodies were detected with biotinylated donkey anti-rabbit (1:500; Jackson ImmunoResearch). Sections were incubated in avidin-biotin complex (1:300, ABC Kit, Vector Lab, PK-6100) and the reaction products were visualized with 3,3’-diaminobenzidine (DAB substrate kit, Vector Lab, SK-4100). Images of Aβ plaques were captured using Keyence BZ-9000 and NPY staining with Zeiss Axio ScanZ.1. Quantification was performed using Image J, with appropriate threshold and particle size settings for Aβ plaques and densitometric measurement for NPY immunoreactivity using as previously described (Palop et al., 2011b).

### Software

The custom software scripts used for raw data processing were written in Perl 5 with modules available on CPAN, and executed on Mac OS X. All software is available upon request and will be provided under GNU GPL licensing.

### Data analysis

Data analysis and statistical testing was performed using GraphPad Prism version 9.3.1. The difference between two data sets was assessed using unpaired (for most experiments) or paired/matched (for data for which the same mouse is both control and treated at different time), two-tailed Student’s t-test. For more than two data sets, differences were assessed by one-way ANOVA (unpaired or paired/matched as appropriate) with correction for multiple-hypothesis testing (Tukey, Šídák, or Holm-Šídák methods as appropriate). Only relevant comparisons were tested. Regression plots show best-fit linear regression lines with 95% confidence interval (CI) for scatterplots of per-minute data. P values for linear regression fits were calculated in Prism using a method equivalent to ANCOVA, first for testing differences in slope and then testing differences in Y-intercept if there was no difference in slope. The method recommendations of Prism were followed unless there was a clear statistical rationale otherwise, and a statistician was consulted when needed. Unless otherwise specified, statistical tests were performed at the level of data resolution presented on a graph -e.g. if a graph shows EDs per minute, statistical tests were performed on minute-level data. All error bars represent SEM unless otherwise specified. Non-significance was defined as a corrected P-value over 0.05. Each figure legend includes specific statistical methods used for the data presented in that figure.

