## Supplemental Figures for "Small-molecule ketone esters treat brain network abnormalities in an Alzheimer’s disease mouse model"

**Newman JC et al.**

**Supplemental Figures 1-10**

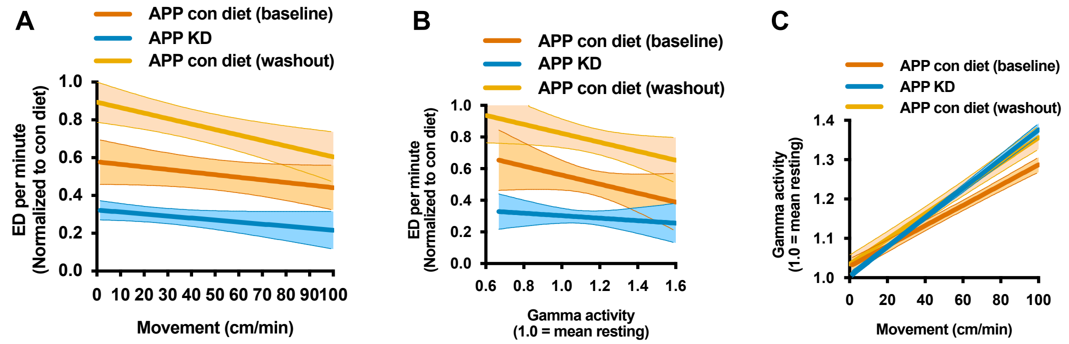


**Supplementary Figure 1**

**Supplementary Figure 1** | Regressions between per-minute movement, ED, and movement-normalized gamma activity, as in Figure 5, for the longitudinal cohort described in Fig. 2A-C. See Fig. 2A for the study schematic. Similar to the other KD experiments, mice fed KD show fewer ED at all levels of movement (**A**) and at all levels of normalized gamma (**B**) as well as no change in the induction of gamma activity by movement (**C**).

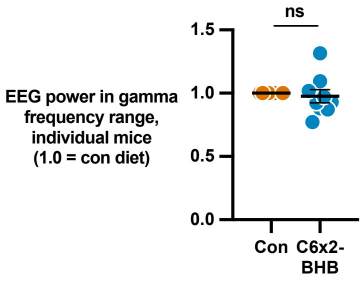


**Supplementary Figure 2**

**Supplementary Figure 2** | The proportion of overall EEG power in the gamma frequency range for individual mice was unchanged by C6x2-BHB. Data are presented as mean ± SEM. P-value via paired T-test.

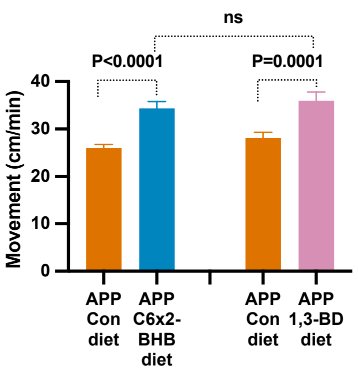


**Supplementary Figure 3**

**Supplementary Figure 3** | Summary of spontaneous movement data during EEGs for the C6x2-BHB and 1,3-butanediol feeding cohorts described in Fig. 4A. Both compounds increased activity modestly, and were not different from each other. Data are presented as mean ± SEM. P-values via one-way ANOVA with Šídák multiple comparisons test.

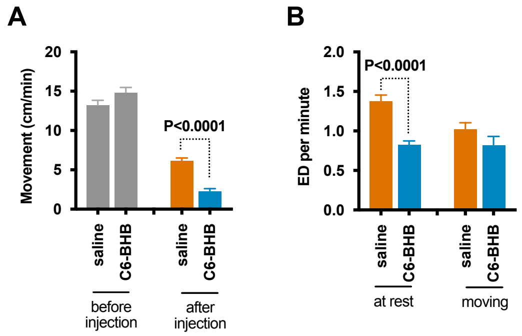


**Supplementary Figure 4**

**Supplementary Figure 4** | Summary of movement data before and after injections in the experiment described in Fig. 3. **A**, C6-BHB reduced movement compared to saline injection. **B**, Stratification of non-normalized ED by movement, averaged per-minute. C6-BHB particularly reduced EDs during otherwise high-spike resting periods. “At rest” is 0 cm movement during the minute. Data are presented as mean ± SEM. P-values via one-way ANOVA with Tukey multiple comparisons test.

A

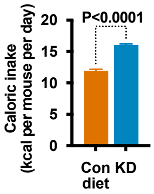


**Supplementary Figure 5**

**Supplementary Figure 5 |** Caloric intake for the cohort described in Fig. 6E-L. Caloric intake is higher for mice fed KD than control diet. Mice were grouped housed with mixed-genotype littermates, so caloric intake cannot be stratified by genotype.

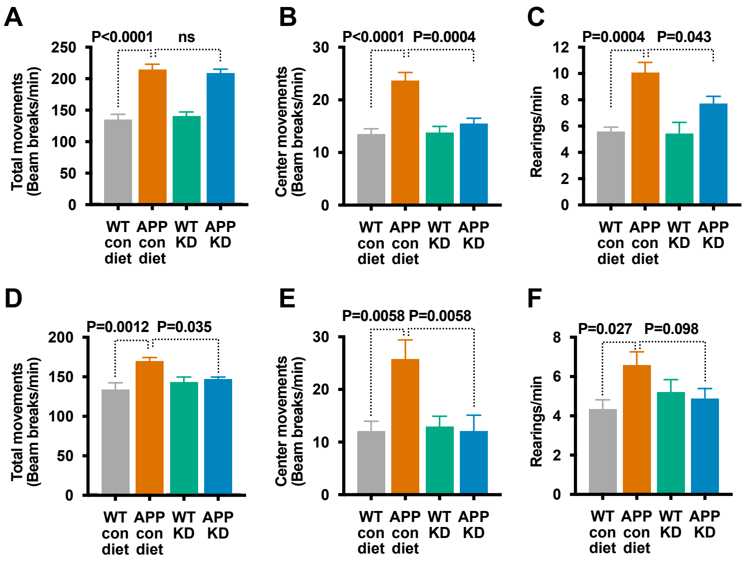


**Supplementary Figure 6**

**Supplementary Figure 6 |** Habituation to the open field data for the cohort described in Fig. 6E-L. As per the schematic in Fig. 6E, mice underwent two habituation test periods from days 15-44 and 128-130. **A-C**,  First habituation to the open field, with training open fields on days 15 and 29 after diet start; data is shown from the test open field on day 44. hAPPJ20 mice on KD have similar overall movement (**A**) but reduced exploratory center movements (**B**) and rearings (**C**) compared to hAPPJ20 mice on control diet indicating partially improved habituation. **D-F**, Second habituation to open field, with training open fields on days 128 and 129 after diet start; data is shown from the test open field on day 130. Compared to hAPPJ20 mice on control diet, hAPPJ20 mice on KD demonstrate improved habituation including reduced total movement (**D**), exploratory center movements (**E**), and rearings (**F**). Data are presented as mean ± SEM. P-values via one-way ANOVA with Šídák multiple comparisons test.

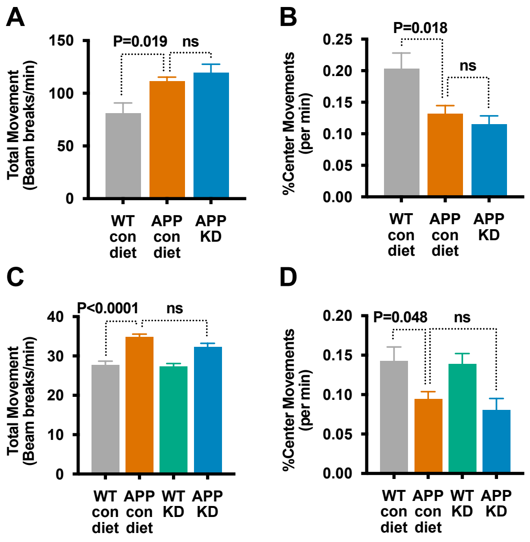


**Supplementary Figure 7**

**Supplementary Figure 7 |** Effect of KD on anxiety phenotypes during the first exposures to the open field. **A-B**, Activity patterns during the first exposure to the open field for the cohort described in Fig. 6A-D, which occurred two weeks after diet start. Compared to WT mice, hAPPJ20 mice show characteristic hyperactivity in the form of increased total movements (**A**), as well as decreased proportion of movement in the center, a finding usually interpreted as increased anxiety. KD does not affect these phenotypes on this first open field (before any habituation can occur). **C-D**, Activity patterns during the first exposure to the open field for the cohort described in Fig. 6E-L show a similar result with hyperactivity (**C**) and increased anxiety for hAPPJ20 mice which is not affected by KD on this initial open field. Data are presented as mean ± SEM. P-values via one-way ANOVA with Šídák multiple comparisons test.

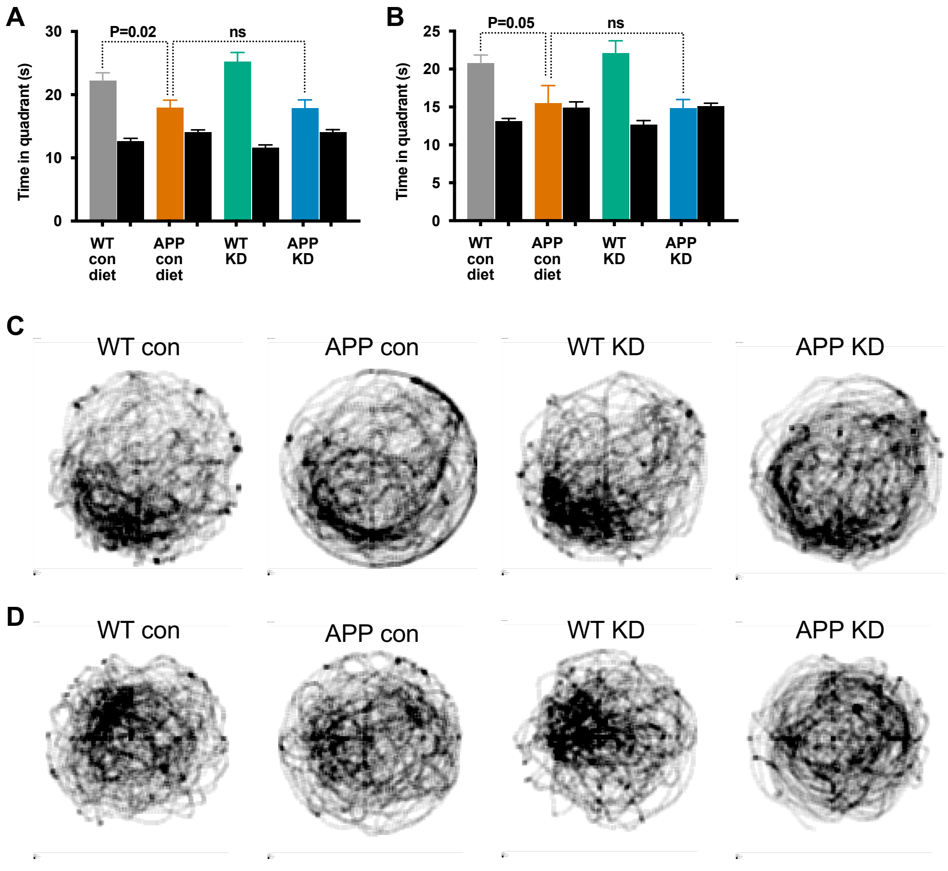


**Supplementary Figure 8**

**Supplementary Figure 8 |** Additional data from Morris water maze in Figure 6I-J. **A-B**, Time spent in the correct target quadrant (colored) versus average of other quadrants (black) during 60 second probe trials following initial training (**A**) and reverse training (**B**). hAPPJ20 mice performed worse than WT, and although KD improved the performance of APP mice on learning trials it did not affect memory performance on these probe trials. **C**, Heat map of search patterns for the probe trial after initial training. Platform had been in the lower-left quadrant. **D**, Heat map of search patterns for the probe trial after reverse training. Platform had been in the upper-left quadrant. Data are presented as mean ± SEM. P-values via one-way ANOVA with Šídák multiple comparisons test.

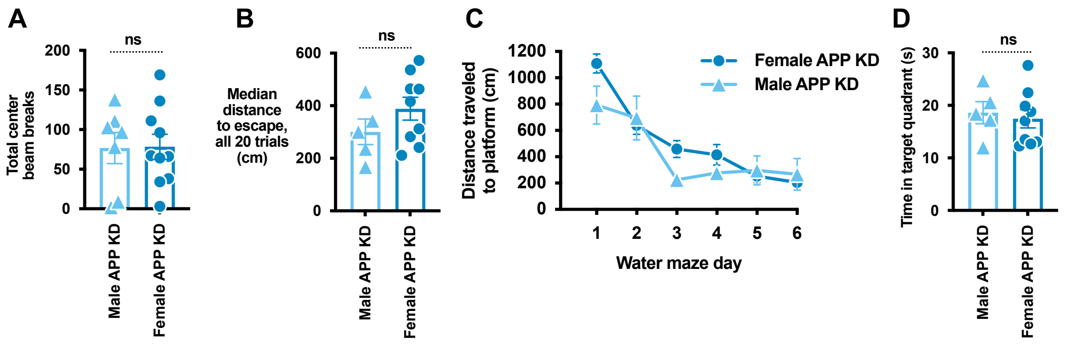


**Supplementary Figure 9**

**Supplementary Figure 9 |** Stratification of learning/memory performance by sex for the cohort described in Fig. 6E-L reveals no significant sex differences. **A**, Performance on the habituation to the open field (represented by total center beam breaks, Fig. 6S2B) stratified by sex. Lower is better. **B**, Overall hidden platform learning trial performance from Fig. 6I stratified by sex. Lower is better. **C**, Daily time course of hidden platform learning trial performance from Fig. 6I stratified by sex. Lower is better. **D**, Probe trial memory performance from Fig. 6S4A stratified by sex. Higher is better. For all panels: Triangles are males; circles are females. Only the APP KD group is shown. Data are presented as mean ± SEM. P-values via unpaired T-test.

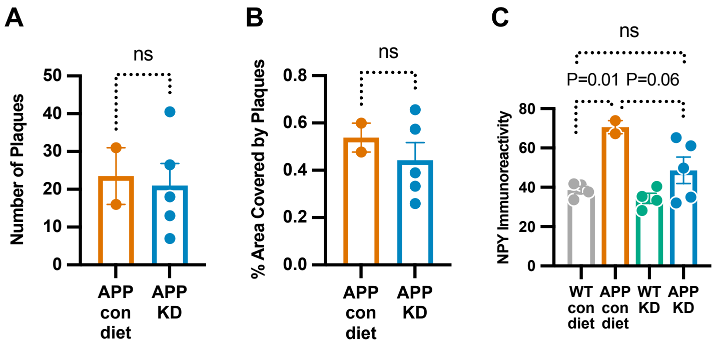


**Supplementary Figure 10**

**Supplementary Figure 10 |** Exploratory immunohistochemical analysis of hAPPJ20 fed KD for 7 months, from the cohort described in Fig. 6E-L. **A-B** Aβ immunohistochemistry. No difference was observed in the number (**A**) or total area (**B**) of Aβ plaques. **C**, Expression of neuropeptide Y (NPY) is increased under conditions of network hyperactivity such as the epileptiform phenotype of hAPPJ20. NPY expression trended lower in KD-fed hAPPJ20, and was not significantly different from WT. Data are presented as mean ± SEM. P-values via unpaired T-test (A, B) or one-way ANOVA with Šídák multiple comparisons test (C).
